# Slow peak alpha frequency and corticomotor depression linked to high pain susceptibility in transition to sustained pain

**DOI:** 10.1101/278598

**Authors:** DA Seminowicz, T Thapa, AJ Furman, SJ Summers, R Cavaleri, JS Fogarty, GZ Steiner, SM Schabrun

**Author notes:** Corresponding author: Dr. Siobhan Schabrun, School of Science and Health, Western Sydney University, Campbelltown Campus, Locked bag 1797, Penrith 2751, NSW, Australia.

## Abstract

Mechanisms that predict an individual’s susceptibility to pain, before pain is present or in the first few days following pain onset, are unknown. We utilised a clinically-relevant human transitional pain model (intramuscular injections of nerve growth factor) to examine brain mechanisms that predict pain susceptibility. Resting state EEG and corticomotor excitability measured by TMS were evaluated longitudinally in healthy individuals as pain developed and resolved over 21 days. Whereas pre-pain central peak alpha frequency (PAF) correlated with peak pain occurring 4-6 days later, altered corticomotor excitability developed several days after pain onset and showed two distinct patterns (facilitation, depression). Individuals with combined slow PAF and corticomotor depression developed more severe pain. These data provide the first evidence of the temporal profile of key brain mechanisms as pain progressively develops. PAF and corticomotor excitability could represent biomarkers for susceptibility to high pain severity and subsequently, the development of chronic pain.

## Introduction

Chronic musculoskeletal pain is a common and debilitating condition (Institute of Medicine, 2011) with few effective treatments. Neuroimaging research has yielded important knowledge about the brain changes associated with the symptoms of chronic pain (Davis & Seminowicz, 2016). A remaining challenge is to identify mechanisms that can predict who will develop chronic pain following an acute pain episode.

Altered brain structure and function can predict the development of chronic low back pain in patients with sub-acute pain (Baliki et al., 2012; Mansour et al., 2013). Yet, mechanisms that can predict patient outcome before pain is present, or in the first few days after pain onset,are unknown. Currently, the best early predictor of chronic pain is high pain severity in response to the intial injury or insult (Walton et al., 2013; Gurcay et al., 2009). However, mechanisms that predispose certain individuals to high pain severity remain elusive. The identification of mechanisms that can detect ‘at risk’ individuals at the time of pain onset would have significant advantages, given that early detection would facilitate early intervention and prevention.

A key limitation in this field has been the absence of a clinically-relevant, human transitional pain model that can be used to investigate pre-pain and early pain mechanisms that can predict who will experience high pain severity, an issue compounded by the inability to obtain pre-pain baseline data on patients with clinical pain. Here, we used a novel human pain model (repeated intramuscular injection of nerve growth factor [NGF]) to induce progressivley developing, sustained muscle pain, lasting up to 21 days (Hayashi, Shiozawa, Ozaki, Mizumura, & Graven-Nielsen, 2013) to examine pre-pain and early-pain (first few days after pain onset) mechanisms of pain susceptibility. We have shown previously that peak alpha frequency (PAF) measured using electroencephalography (EEG) during a pain-free period correlates with pain severity experienced in a capsaicin heat pain model 45 minutes later (Furman et al., 2017). We have also shown, using an identical transitional pain model, that corticomotor excitability and organisation (measured using transcranial magnetic stimulation [TMS]) is altered 4 days after pain onset, (Schabrun, Christensen, Mrachacz-Kersting, & Graven-Nielsen, 2016). In the present study we examined whether these brain mechanisms were related to an individual’s pain susceptibility.

## Results

In Study 1, twenty healthy individuals (11M, 9F) participated in 5 laboratory sessions (Day 0, 2, 4, 6, and 14) and completed pain diaries for 21 days. Nerve growth factor (NGF) was injected into the muscle belly of ECRB on days 0, 2, and 4, causing progressively developing, sustained muscle pain (Schabrun et al., 2016). At each session, eyes closed resting state electroencephalography (EEG), transcranial magnetic stimulation (TMS) motor cortex mapping methods and pressure pain thresholds (PPTs) were recorded. Individuals were divided into two groups based on whether they showed facilitation (n=8) or depression (n=12) of corticomotor excitability in response to pain (Fig 1A). Three individuals did not develop pain and were excluded. To further explore EEG effects, an independent group of 11 participants (5M, 6F) underwent similar procedures with NGF injections on days 0 and 2, and measurements up to day 4 (Study 2).

**Fig 1.**
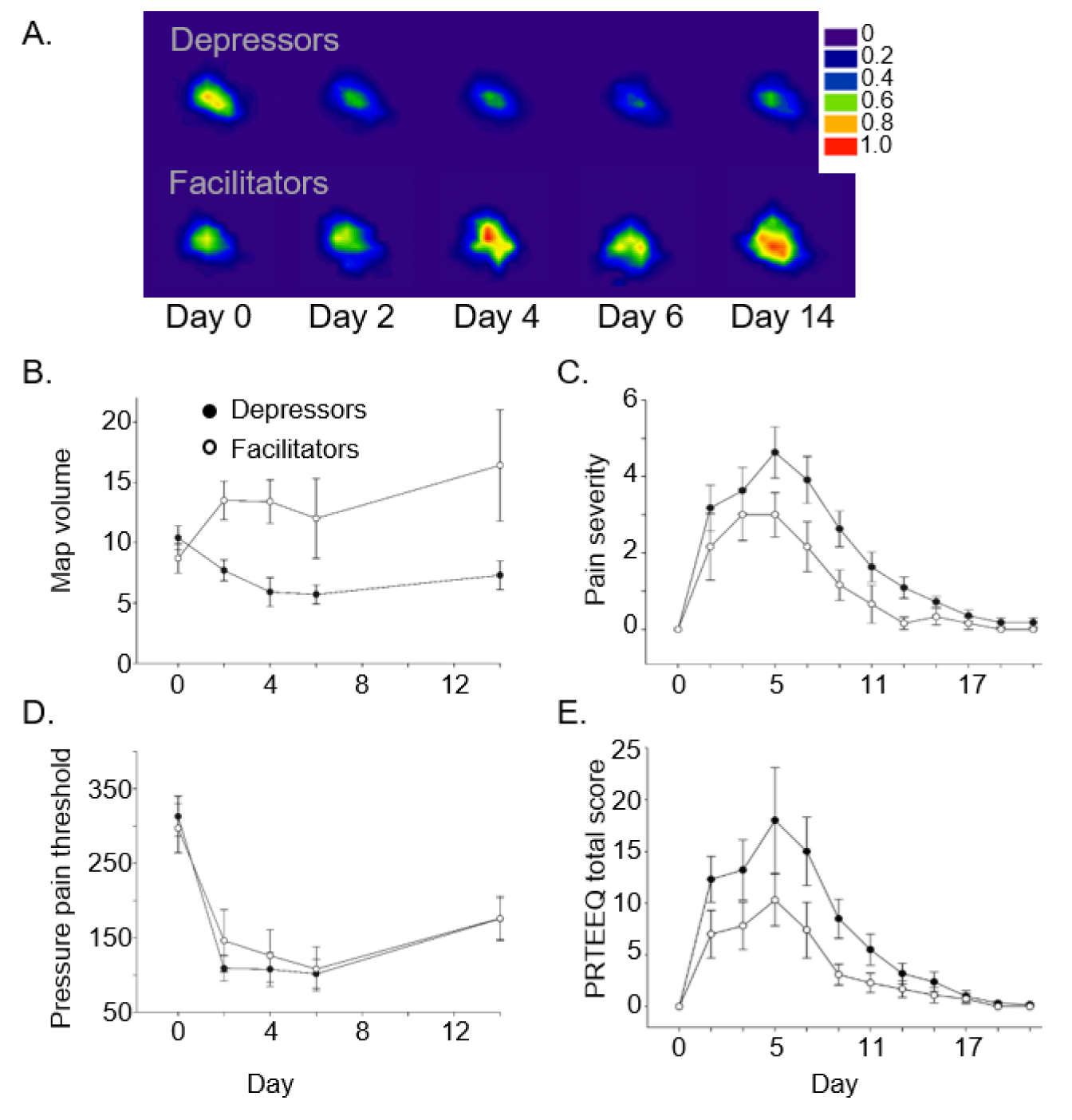
Corticomotor excitability is associated with pain severity. **A.** Averaged motor cortex maps for the ECRB muscle in depressors (n=11, top) and facilitators (n=6, bottom) in response to NGF-induced sustained muscle pain (study 1, n=20). Maps are normalized to Day 0 for each participant. Warmer colours represent greater excitability (proportion of the motor evoked potential response at Day 0, mV). **B-E.** Group data (mean±standard error) in depressors (filled circles) and facilitators (open circles) for: **B.** motor cortical map volume; **C.** pain severity (numerical rating scale); **D.** pressure pain thresholds (kPa); **E.** disability (patient rated tennis elbow evaluation questionnaire total scores, PRTEEQ). Depressors developed more pain than facilitators but did not display greater sensitivity to pressure.

Change in corticomotor excitability was significantly different between facilitator and depressor groups (F_1,15_=6.6, p<0.05; Fig 1B). There were no differences between groups for age, depression, anxiety, or pain catastrophizing (all p>0.1) and when these factors were included as covariates, the difference in corticomotor excitability remained (F_1,11_=5.1, p<0.05). Pain was greater in the depressor group (F_1,15_=5.1, p<0.05) and disability showed a similar, non-significant trend (F_1,15_=2.8, p=0.11; Fig1C,E). These unique data reveal two distinct patterns of motor plasticity in response to pain (facilitation or depression) that are not explained by demographic or psychosocial factors. Corticomotor facilitation in response to sustained pain is hypothesized to reflect an adaptive motor learning response that enhances the search for a new movement strategy to maximise task performance while minimising pain (Schabrun et al., 2016). This theory is consistent with our observation that individuals who exhibited corticomotor facilitation had less pain than those who exhibited depression.

PPTs at the site of injection, but not a remote site (tibialis anterior), decreased following NGF injection in both groups (F_1,15_=21.6, p<.001; Fig 1D). Intramuscular injection of NGF is known to sensitize high threshold mechanosensitive afferents via direct effects on trkA-expressing nociceptors or indirectly via mast cells, sympathetic postganglionic neurons or accumulation of neurotrophins at the site of injection (Pezet & McMahon, 2006). Although depressors developed significantly more pain, PPTs decreased to the same extent as facilitators (F_1,15_=0.08, p>.5), suggesting the degree of peripheral sensitization was similar between groups. This finding implies that an individual’s susceptibility to pain is mediated by central mechanisms.

We next examined the relationship between baseline resting state peak alpha frequency (PAF) and pain severity. We have shown that PAF is inversely correlated with pain severity occurring after 45 minutes of capsaicin-induced heat pain (Furman et al., 2017), consistent with findings in neuropathic pain where chronicity is associated with slowed PAF (Sarnthein, Stern, Aufenberg, Rousson, & Jeanmonod, 2006; de et al., 2013a). Remarkably, in the NGF model, pre-pain PAF was negatively correlated with peak pain occurring 4–6 days later, suggesting that baseline PAF could be a biomarker for pain severity before pain develops (Fig 2). Indeed, when individuals were divided into four groups based on PAF (fast (mean PAF±SD; 10.04±0.08) or slow (9.80±0.05) defined using median split) and corticomotor excitability (facilitation or depression; Fig 3A), individuals with slow PAF and corticomotor depression had greater peak pain than any other group (F_3,27_=3.0, p<0.05; Fig 3B), suggesting this combination of cortical mechanisms is related to an individual’s pain susceptibility.

**Fig 2.**
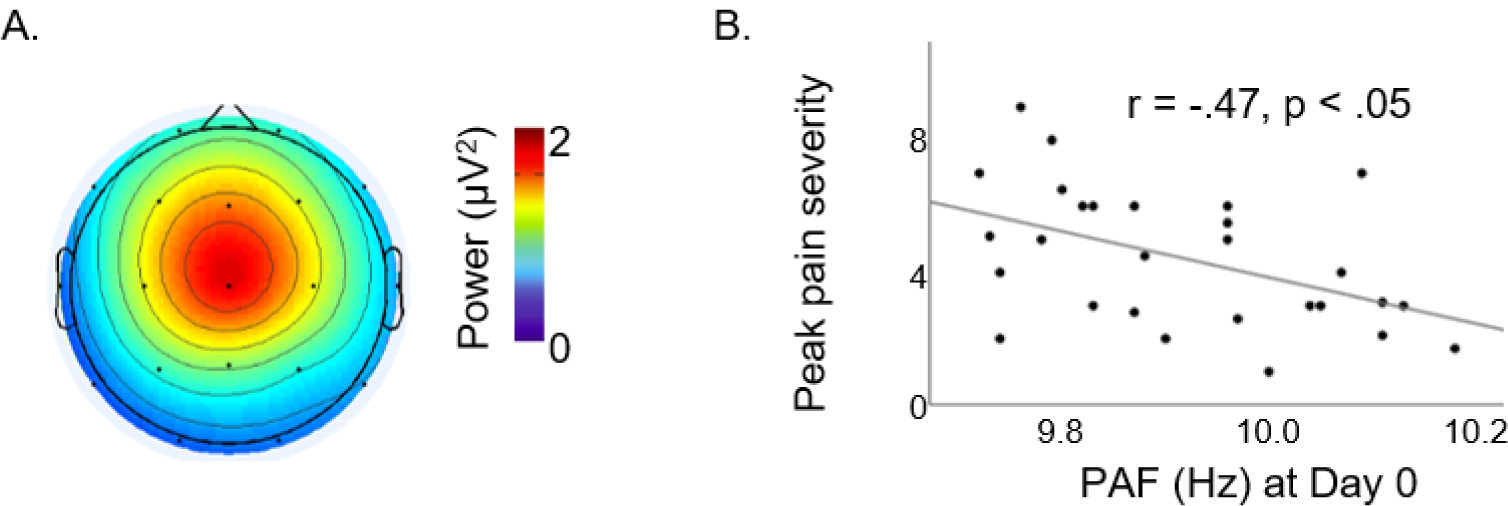
Pre-pain Central Peak Alpha Frequency (PAF) is associated with pain severity. Eyes closed resting state central PAF recorded at baseline, pain-free state, is inversely correlated with peak pain severity from the NGF model, occurring 4–6 days later (studies 1 and 2, n=28).**A.** group averaged component map from ICA method used to identify central PAF component for each participant. **B.** correlation between center of gravity PAF at baseline with peak pain severity.

**Fig 3.**
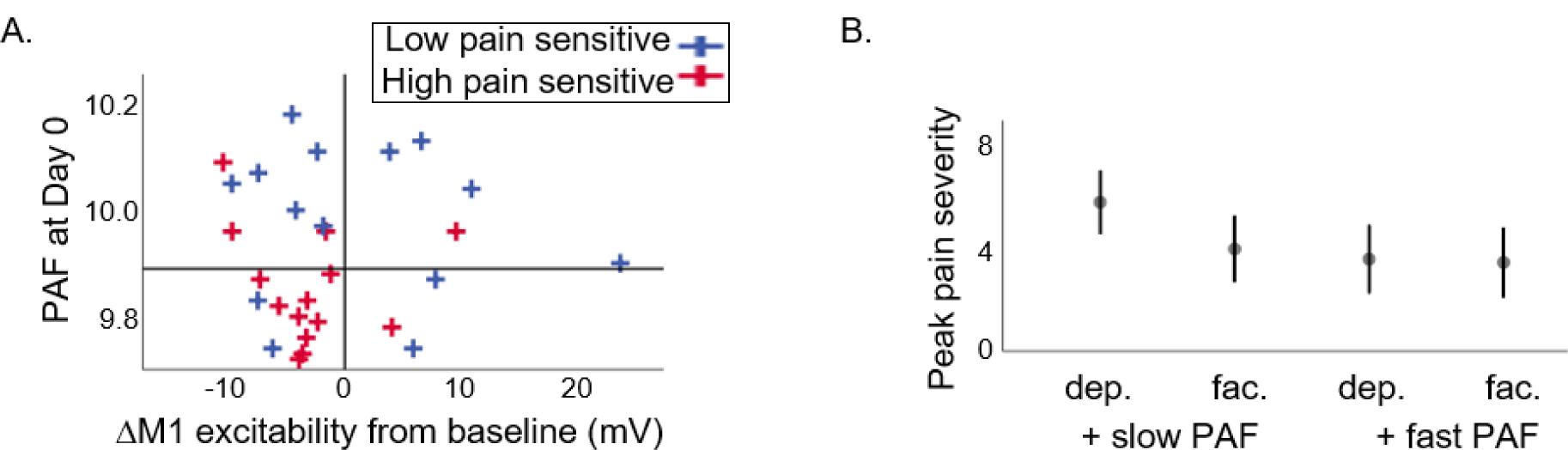
Combined slow PAF and corticomotor depression is associated with pain susceptibility. Combination of slow baseline PAF and corticomotor depression (studies 1 and 2, n=28). **A.** Quadrant plot shows the median split PAF and depressor/facilitator split for individuals who are high and low pain senstive based on median split of pain severity ratings. High pain sensitive individuals cluster in the bottom left quadrant, representing depressors with slow PAF. **B.** Slow PAF/depressor group has significantly higher pain than other groups. Mean ± SEM of peak pain recorded between days 4-6.

## Discussion

In a clinically-relevant model of sustained muscle pain, we demonstrate that coricomotor depression is associated with higher pain severity than corticomotor facilitation. A similar trend was observed for disability. However, there were no differences between facilitators and depressors in pressure pain thresholds of the ECRB, which dramatically decreased following NGF injection, suggesting differences in pain susceptibility are centrally mediated. We further showed that EEG PAF recorded pre-pain was correlated with peak pain severity occurring 4 to 6 days later. Notably, individuals with combined slow PAF and corticomotor depression had greater pain severity than individuals with any other combination of mechanisms. In other words, these two factors appear to predict pain susceptibility. The combination of slow PAF and corticomotor depression is a plausible biomarker for the development of high pain severity. Our findings are important given that high pain severity is a known risk factor for the development of chronic musculoskeletal pain (Walton et al., 2013; Gurcay et al., 2009).

Numerous studies have suggested a link between altered corticomotor excitability and the symptoms of chronic musculoskeletal pain (Schabrun, Elgueta-Cancino, & Hodges, 2017; Tsao, Galea, & Hodges, 2008; Te, Baptista, Chipchase, & Schabrun, 2017; Tsao, Danneels, & Hodges, 2011; Schabrun, Hodges, Vicenzino, Jones, & Chipchase, 2015). However, only one study has conducted a longitudinal investigation of this mechanism in response to pain. Using the NGF pain model, that study demonstrated increased corticomotor excitability at the group level 4 days following pain onset (Schabrun et al., 2016). Consistent with recent theories on pain adaptation, increased corticomotor excitability is interpreted to reflect the search for a new motor strategy that utilizes synergies with surrounding muscles to redistribute muscle activity and reduce loading on a painful structure {Masse-Alarie, 2017 11035 /id;Hodges and Tucker 2011). This hypothesis is further supported by observations of increased corticomotor excitability in the early stages of motor learning (Pascual-Leone et al. 1994; Pascual-Leone et al. 1995) and by observations of increased movement variability in the acute stage of pain (Hodges et al. 2013). Our data extend these findings. When individual-level data are considered, two distinct patterns of corticomotor excitability are observed: facilitation or depression. Individuals who demonstrate facilitation have lower pain, supporting the hypothesis that increased corticomotor excitability is an adaptive cortical response when pain is present. Conversley, corticomotor depression is associated with higher pain, and in combination with pre-pain PAF, these mechanisms appear to predict individuals at risk of high pain severity.

Alpha (8–12Hz) is the predominant oscillatory activity in the EEG signal. Peak alpha frequency recorded during an eyes closed resting state is one alpha metric related to the pain experience. Resting state PAF varies considerably amongst individuals (Bazanova & Vernon, 2014), but it is stable in test-retest studies (Grandy et al., 2013) and may be a heritable phenotypic trait (Posthuma, Neale, Boomsma, & de Geus, 2001; Smit, Wright, Hansell, Geffen, & Martin, 2006). In chronic pain patients, alpha rhythms are slower in frequency and elevated in power relative to matched, healthy controls suggesting that PAF alterations may represent factors such as disease progression or pain vulnerability (Sarnthein et al., 2006; de et al., 2013b; Lim, Kim, Kim, & Chung, 2016; Walton, Dubois, & Llinβs, 2010). Our previous findings indicate that PAF can predict future experiences of pain: we showed a relationship between alpha and pain sensitivity in 21 healthy subjects (Furman et al., 2017). The mechanistic link between PAF and pain is still unclear, but emerging evidence from other sensory domains have begun to provide important clues. In the visual and auditory domains, PAF has been shown to play a role in determining the rate of sensory sampling with faster PAFs associated with more fine-grained temporal resolution {Cecere, 2015 10971 /id;Minami, 2017 11051 /id}. Alongside electrophysiological findings that neuronal excitability is modulated by the phase of the alpha rhythm in the thalamus (Lorincz, Kekesi, Juhasz, Crunelli, & Hughes, 2009) or sensorimotor cortex (Haegens, Nacher, Luna, Romo, & Jensen, 2011), these results suggest that PAF plays an important role in determining how many times per second information about the external environment is obtained and relayed through neuronal circuits. Although it is not immediately clear to what extent or why fast and slow PAF are associated with M1 facilitation and depression, respectively, PAF may play a role in shaping cortical responses to a noxious input. Although the link between Alpha power and BOLD activations across the cortex are well characterized (Mayhew & Bagshaw, 2017), the impact of PAF on this relationship is currently untested. Future EEG-fMRI studies investigating whether variations in PAF shape pattern of cortical activity in the presence and absence of pain may provide important insight into why PAF is such a strong, reliable predictor of future pain.

In summary, we have shown that two mechanimisms – peak alpha frequency and corticomotor excitability – are related to future pain severity in a long-lasting muscle pain model in otherwise healthy participants. Individuals with slow PAF and corticomotor depression had higher pain than all other groups. This phenotype could represent a risk marker for the development of chronic pain following injury or surgery.

## Materials and Methods

### Participants

Thirty-one healthy, right-handed individuals participated (Study 1 included 20 individuals: 11 male; 23±4.3 years; mean±standard deviation [SD]; Study 2 included 11 individuals: 5 male; 24±6.9 years). Handedness was assessed using the Edinburgh handedness questionnaire (Oldfield, 1971). Participants with a history of neurological, psychiatric, musculoskeletal or upper limb conditions were excluded and a transcranial magnetic stimulation (TMS) safety screen was completed prior to study enrolment(Keel, Smith, & Wassermann, 2001). All participants provided written informed consent consistent with the Declaration of Helsinki. Experimental procedures were approved by the institutional ethics committee (H10184).

### Experimental protocol

Study 1. Participants attended the laboratory on 5 occasions: Days 0, 2, 4, 6, and 14 (Fig 1). Day 0 (pre-pain) outcome measures included eyes closed resting state electroencephalography (EEG), pressure pain thresholds (PPTs) and TMS-derived motor cortical maps. The State-Trait Anxiety Inventory(Spielberger, Gorsuch, & Lushene, 1970), the Beck Depression Inventory (BDI)(Beck, Ward, Mendelson, Mock, & Erbaugh, 1961) and the Pain Catastrophizing Scale (PCS)(Sullivan, Bishop, & Pivik, 1995) were also administered on Day 0. Assessment of PPTs and motor cortical maps were repeated on Days 2, 4, 6 and 14. Nerve growth factor (NGF) was injected into the belly of the right extensor carpi radialis brevis (ECRB) muscle immediately following collection of all outcome measures on Day 0, 2 and 4. No injection was given on Day 6 or 14. Electronic pain diaries were administered on each alternate day from Day 1 to Day 21 (Day 1,3, 5…21).

Study 2. Participants attended the laboratory on 3 occasions: Days 0, 2 and 4 (Fig 1). Day 0 (pre-pain) outcome measures included eyes closed resting state EEG, PPTs and TMS-derived motor cortical maps. Assessment of PPTs and motor cortical maps were repeated on Days 2 and 4. NGF was injected into the belly of the right ECRB immediately following collection of all outcome measures on Days 0 and 2. Electronic pain diaries were administered on Days 0, 2 and 4.

### NGF-induced muscle pain

Nerve growth factor, an endogenous neuromodulator, is essential for the development and maintenance of the nervous system (Lewin, Ritter, & Mendell, 1992). Intramuscular injection of NGF in humans has been shown to induce progressively developing, clinically-relevant muscle pain that is sustained for up to 21 days and is accompanied by muscle hyperalgesia, movement-evoked pain and reduced function (Bergin et al., 2015). After cleaning the skin with alcohol, a dose of 5 μg (0.2 ml) sterile, recombinant human NGF was given as a bolus injection into the muscle belly of ECRB using a 1 ml syringe with a disposable needle (27G)(Schabrun et al., 2016).

### Electronic pain diary

Pain was assessed using an 11-point numerical rating scale (NRS) anchored with ‘no pain’ at zero and ‘worst pain imaginable’ at 10. The Patient-rated Tennis Elbow Evaluation Questionnaire (PRTEEQ) was used to assess disability(Macdermid, 2005). Scores for pain (sum of five items out of a maximum score of 50) and disability (sum of ten items, divided by 2, out of a maximum score of 50) were combined to give a total score ranging from 0 (no pain and no functional impairment) to 100 (worst pain imaginable with significant functional impairment).

### Resting state electroencephalography

EEG assessments were the first procedures performed at each laboratory session in both Studies 1 and 2. EEG data were recorded continuously DC–70 Hz from 19 scalp sites (Fp1, Fp2, F3, Fz, F4, F7, F8, C3, Cz, C4, P3, Pz, P4, P7, P8, T7, T8, O1, and O2) and M2 with an electrode cap using sintered Ag/AgCl electrodes. M1 was used as the active reference and the cap was grounded by an electrode located between AF3 and AF4. Electro-oculogram (EOG) was recorded using sintered Ag/AgCl electrodes placed 2 cm above and below the left eye for vertical movements, and on the outer canthus of each eye for horizontal movements. Data were acquired at 1000 Hz with the default gain setting and a 50 Hz notch filter using a Neuroscan Synamps 2 digital signal-processing system and Neuroscan 4.3.1 Acquire software. Impedances were < 5 kΩ for cap, EOG, and reference electrodes. Six minutes of continuous resting state EEG was recorded at the beginning of the session. Participants were asked to sit quietly while recordings of eyes-open followed by eyes-closed resting states were acquired for three minutes each; eyes closed data were used for all analyses.

### EEG Data Processing

The primary data of interest were the eyes closed resting state EEG data collected prior to any NGF injection on Day 0. Data were re-referenced to digitally linked mastoids (i.e., M1 & M2) using Neuroscan 4.5 Edit software. Preprocessing of EEG data was performed in EEGLAB 13.6.5b (Delorme & Makeig, 2004) using an approach similar to that used previously (Furman et al., 2017; Scheeringa, Mazaheri, Bojak, Norris, & Kleinschmidt, 2011). First, continuous EEG data were filtered between 2 and 16 Hz using a linear FIR filter. Next,extended Infomax independent component analysis (ICA) was performed to decompose the eyes closed resting state data to a series of 19 spatial components.

### Quantification of PAF

A Discrete Fourier transformation (DFT) was done on the time series of each component to obtain a frequency-power spectra for each of the 19 spatial components. The frequency decomposition of the component data was performed using routines in FieldTrip (Oostenveld, Fries, Maris, & Schoffelen, 2011). The data were segmented into 5 second epochs and power spectral density in the 2–16 Hz range was derived for each epoch in 0.2 Hz bins. A Hanning taper was applied to the data prior to spectra calculation to reduce edge artifacts (Mazaheri et al., 2010; Mazaheri, Nieuwenhuis, van, & Jensen, 2009; Mazaheri et al., 2014).

For each participant, we visually inspected the frequency-spectra of the components, and identified components that had both a clear alpha peak (7–14 Hz; a wider range was included to ensure individual variability in alpha peak frequency was detected) and a projected scalp topography over the Cz sensor that resembled the “sensorimotor” component template generated from our previous study (Furman et al., 2017). In cases where multiple components met these criteria, we selected the spatial component with the highest variance contribution.

The peak alpha frequency for each 5 second epoch was estimated using a center of gravity (CoG) method (Jann et al., 2012; Jann, Koenig, Dierks, Boesch, & Federspiel, 2010; Klimesch, Schimke, & Pfurtscheller, 1993). We defined CoG as follows:

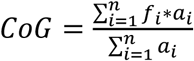

where *f*_*i*_ is the *i*th frequency bin including and above 9 Hz, *n* is the number of frequency bins between 9 and 11 Hz, and *a*_*i*_ the spectral amplitude for *f*_*i*_. PAF were estimated for the selected “sensorimotor” alpha components for every 5 second epoch and then averaged for each subject. From our previous work, we have determined that this narrow analysis band reduces the influence of 1/*f* EEG noise on the estimation of PAF (Furman et al., 2017).

### Pressure algometry

Pressure was applied perpendicular to the surface of the skin using a hand-held pressure algometer with a 1cm^2^ probe (Force Ten, FDX Force Gage, Wagner instruments, Greenwich, USA). Three readings at the pressure pain threshold were made at 1-min intervals at three sites:i) right ECRB (injected site), ii) left ECRB (matched to site of injection on right ECRB), and iii) left tibialis anterior (10 cm distal from the tibial tuberosity). A tape measure was used to measure the position of each site (ECRB – distance (cm) distal to the lateral epicondyle and medial distance (cm); Tibialis Anterior – distance (cm) distal from the base of the patellar tendon) and these values were recorded to ensure consistent positioning across measurement sessions. Participants were instructed to verbalize when the sensation of pressure first changed to pain. The average of the three recordings at each site was used for analysis.

### Motor cortical maps

Electromyographic (EMG) activity was recorded from right ECRB using bipolar Ag/AgCl surface electrodes (Medicotest 720-01-K, Ambu A/S, Ballerup, Denmark) positioned over the muscle belly. EMG signals were amplified (1000 times), bandpass filtered between 20-1000 Hz and sampled at 2 kHz (CED 1401 AD, Cambridge Electronic Design, Cambridge, United Kingdom) using Signal acquisition software (CED, version 5.08 x 86).

A standard procedure for mapping the motor cortical representation of upper limb muscles was used (Schabrun et al., 2016; Schabrun et al., 2015; Schabrun, Stinear, Byblow, & Ridding, 2009; Schabrun & Ridding, 2007). Participants were fitted with a cap marked with a 1 cm x 1 cm grid and orientated to the vertex (point 0,0). A standard 70 mm figure-of-eight coil connected to a magnetic stimulator (Magstim 200, Magstim Co. Ltd. Dyfed, UK) was used to provide single-pulse TMS. The coil was positioned tangentially to the scalp with the handle pointing posterolaterally at a 45° angle from the mid-sagittal plane. This orientation is optimal for the induction of posterior-to-anterior (PA) directed current for trans-synaptic activation of horizontal cortical connections in M1 (Bashir, Perez, Horvath, & Pascual-Leone, 2013; Brasil-Neto et al., 1992). The optimal site (‘hotspot’) for eliciting motor evoked potential (MEP) responses from the relaxed ECRB muscle was determined by systematically moving the coil in 1 cm increments around the motor cortex. The site that evoked the largest responses at the lowest stimulator intensity was considered the hotspot. The stimulus intensity for mapping was set at 120% of resting motor threshold (rMT), defined as the minimum stimulator intensity at which 5 out of 10 stimuli applied at the hotspot evoked a response with a peak-to-peak amplitude of at least 50 μV. This intensity was determined on Day 0 and kept constant for mapping on Days 2, 4, 6 and 14. Single-pulse TMS was applied every 6 seconds with a total of 5 stimuli at each site. The number of scalp sites was pseudorandomly increased until no MEP was recorded (defined as less than 50 μV peak-to-peak amplitude in all five trials in all border sites)(Schabrun et al., 2015; Schabrun & Ridding, 2007). Participants were seated and instructed to keep their hand and forearm relaxed with the wrist pronated throughout the experiment. All TMS procedures adhered to the TMS checklist for methodological quality(Chipchase et al., 2012).

The number of active map sites and map volume was calculated. A site was considered ‘active’ if the mean peak-to-peak amplitude of the five MEPs evoked at that site was greater than 50 μV. The mean peak-to-peak MEP amplitudes at all active sites were summed to calculate the map volume.

### Statistical analyses

Participants were divided into two groups based on whether they showed facilitation or depression of corticomotor excitability in response to injection of NGF (average map volume across days 2, 4, 6 and 14 as a proportion of baseline). We performed a mixed model ANOVA for each dependent variable of interest for the between-subjects factor of group (facilitator, depressor) and the within-subjects factor of time (Days 1, 3, 5,….21 for pain and disability;Days 0, 2, 4, 6 and 14 for corticomotor excitability and PPTs) in Study 1, which was designed to examine the central mechanisms of a long-lasting pain model. PAF was analysed for the whole study sample (Studies 1 and 2, that had identical methods up to day 4). Peak pain severity for each individual was correlated with baseline PAF using Pearson correlation. To examine the interaction of corticomotor excitability and PAF, we created four groups via median split (slow/fast PAF by depressor/facilitator) and compared peak pain severity using one-way ANOVA.

---

**Fig S1:**
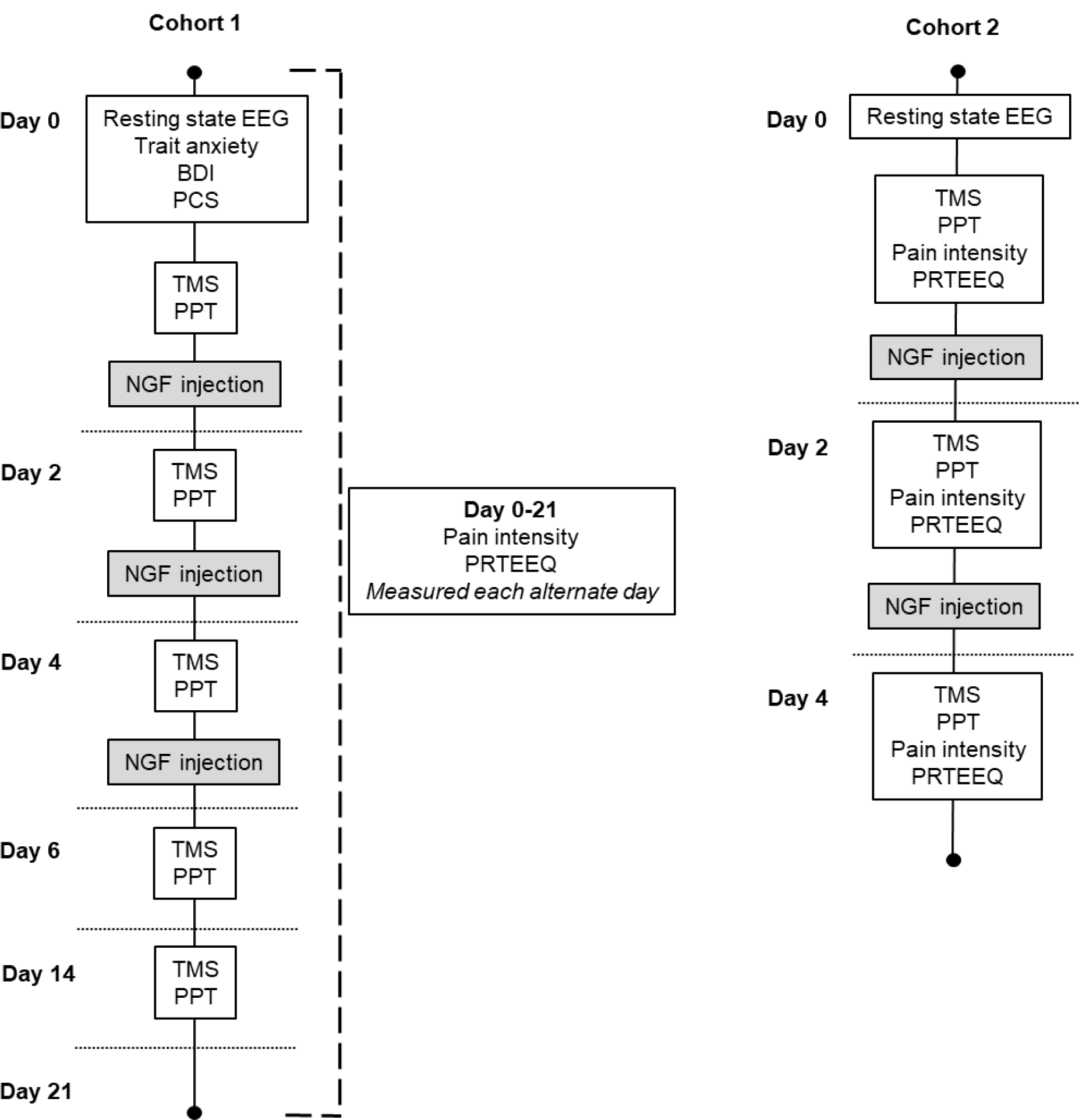
Experimental protocol for cohort 1 (left panel) and cohort 2 (right panel). EEG – electroencephalography, BDI – Beck Depression Inventory, PCS - Pain Catastrophizing Scale, TMS – Transcranial magnetic stimulation, PPT – Pressure pain threshold, NGF–Nerve growth factor, PRTEEQ – Patient Rated Tennis Elbow Evaluation Questionnaire.

## Acknowledgements

SMS receives salary support from The National Health and Medical Research Council (NHMRC) of Australia (#1105040). GZS receives salary support from a NHMRC-Australian Research Council Dementia Research Development Fellowship (#1102532). DAS received an International Visiting Research Fellowship from Western Sydney University. SJS and RC hold Australian Postgraduate Awards. The authors have no competing interests to declare.

